# Revisiting *Leishmania* GP63 host cell targets reveals a limited spectrum of substrates

**DOI:** 10.1101/2022.06.06.494968

**Authors:** Marie-Michèle Guay-Vincent, Christine Matte, Anne-Marie Berthiaume, Martin Olivier, Maritza Jaramillo, Albert Descoteaux

**Affiliations:** Institut national de la recherche scientifique, Centre Armand-Frappier Santé Biotechnologie, Laval, QC, Canada; Department of Microbiology and Immunology, McGill University, Montréal, QC, Canada; Infectious Diseases and Immunity in Global Health Program, The Research Institute of the McGill University Health Centre, Montréal, QC, Canada

## Abstract

Colonization of host phagocytic cells by *Leishmania* metacyclic promastigotes involves several parasite effectors, including the zinc-dependent metalloprotease GP63. The major mode of action of this virulence factor entails the cleavage/degradation of host cell proteins. Given the potent proteolytic activity of GP63, identification of its substrates requires the adequate preparation of cell lysates to prevent artefactual degradation during cell processing. In the present study, we re-examined the cleavage/degradation of reported GP63 substrates when GP63 activity was efficiently neutralized during the preparation of cell lysates. To this end, we infected bone marrow-derived macrophages with either wild type, Δ*gp63*, and Δ*gp63*+*GP63 L. major* metacyclic promastigotes for various time points. We prepared cell lysates in the absence or presence of the zinc-metalloprotease inhibitor 1,10-phenanthroline and examined the levels and integrity of ten previously reported host cell GP63 substrates. Inhibition of GP63 activity with 1,10-phenanthroline during the processing of macrophages prevented the cleavage/degradation of several previously described GP63 targets, including PTP-PEST, mTOR, p65RelA, c-Jun, VAMP3, and NLRP3. Conversely, we confirmed that SHP-1, Synaptotagmin XI, VAMP8, and Syntaxin-5 are *bona fide* GP63 substrates. These results point to the importance of efficiently inhibiting GP63 activity during the preparation of *Leishmania*-infected host cell lysates. In addition, our results indicate that the role of GP63 in *Leishmania* pathogenesis must be re-evaluated.

**AUTHOR’S SUMMARY:** In the protozoan parasite *Leishmania*, the abundant zinc-dependent metalloprotease GP63 is expressed at high levels at the surface of the promastigotes forms of the parasite. Upon phagocytosis by host macrophages, this metalloprotease is released from the parasite’s surface and spreads across the cytosol of infected cells. There, GP63 cleaves a number of host cell proteins involved in the control of host microbicidal function and in the regulation of immune responses, thereby contributing the ability of *Leishmania* to impair host defence mechanisms against infection. Given the abundance and powerful proteolytic activity of GP63, it is crucial to prevent artefactual proteolysis during processing of infected cells to identify genuine GP63 substrates. In this study, we found that inhibition of GP63 activity with 1,10-phenanthroline during the processing of macrophages prevented the degradation of several of previously identified GP63 substrates. These results uncover the importance of efficiently inhibiting GP63 activity during the preparation of *Leishmania*-infected host cell lysates.

## INTRODUCTION

The protozoan parasite *Leishmania* is the causative agent of a spectrum of human diseases called leishmaniases (1). Infectious metacyclic promastigotes are inoculated into their mammalian hosts by phlebotomine sand flies (2) and are taken up by host phagocytes, where they differentiate and replicate as mammalian-stage amastigotes (3). During the host cell colonization process, internalized promastigotes inhibit phagolysosome biogenesis, create a specialized parasitophorous vacuole, and evade microbicidal and immune processes by altering key cellular pathways (4-6). *Leishmania* uses a panoply of pathogenicity factors to subvert the antimicrobial activities of host macrophages and to dampen host immune responses (6-11). Among those, the zinc-dependent metalloprotease GP63 has been extensively studied owing to its abundance and role in virulence (6, 7, 12-14). Hence, GP63 was shown to disrupt cellular processes and pathways involved in host defense against infection through the proteolysis of a variety of host cell proteins that include components of signaling cascades, transcription factors, nucleoporins, cytoskeletal proteins, and regulators of membrane fusion (7, 15-17). Collectively, these findings contributed to our understanding of the molecular, cellular, and immune bases of *Leishmania* pathogenesis.

The wide range of host cell proteins identified as GP63 targets are associated to diverse intracellular compartments (7, 15, 17), implying that this parasite protease efficiently accesses different host cell locations to exert its action. Two distinct mechanisms were described for GP63 to reach its targets. One is based on the finding that GP63 is present in extracellular vesicles/exosomes released by promastigotes (18, 19). These exosomes modulate the activity of macrophage protein tyrosine phosphatases (PTPs) and transcription factors, as well as the expression of a number of pro-inflammatory genes (18-20). Whereas the exact mechanism by which GP63 present in these exosomes reaches its targets remains to be identified, it has been proposed that these vesicles may fuse with macrophages once in the phagolysosomal compartment to release their content into the cytoplasm of infected cells (18). In this regard, entry of GP63 was shown to be in part mediated by a lipid raft-dependent mechanism (21). In addition, the bulk of GP63 rapidly sheds from the surface of internalized promastigotes, exits the parasitophorous vacuole through a mechanism involving Sec22b and the secretory pathway, and traffics within the host cell mainly in the endoplasmic reticulum (ER) and the ER-to-Golgi intermediate compartment (22). Disruption of the trafficking between the ER and the Golgi impairs the exit of GP63 from the parasitophorous vacuole and prevents the cleavage of select host cell proteins (22). These observations strongly support the notion that GP63 must traffic within infected cells and access various intracellular compartments to cleave its targets.

The proteolytic activity of GP63 was serendipitously discovered when parasite fractions containing this protein were mixed with molecular weight standards for SDS-PAGE (23, 24). Characterization of the nature of this protease revealed that whereas its activity was not affected by serine or cysteine protease inhibitors, EDTA, or EGTA, it was completely abrogated by the zinc chelator 1,10-phenanthroline, suggesting that GP63 is a zinc metalloprotease (23). Subsequent analysis of the metal content of GP63 by atomic emission and absorption spectroscopy and the presence of a highly conserved zinc binding domain confirmed that GP63 is a zinc metalloprotease (25). Given the abundance and powerful proteolytic activity of GP63, unambiguous identification of its substrates in host cells thus hinges on an adequate preparation of cell lysates to prevent artefactual post-cell lysis proteolysis. However, in a number of previous studies aimed at investigating the impact of GP63 on the cleavage/degradation of host cell substrates, the presence of zinc metalloprotease inhibitors in lysis buffers was not specifically mentioned. To discard the possibility that some host cell proteins were wrongly identified as GP63 substrates due to post-cell lysis proteolytic events, we sought to re-evaluate the cleavage/degradation of previously reported GP63 substrates. In the present study, we show that inhibition of GP63 activity with the metalloprotease inhibitor 1,10-phenanthroline during the lysis of infected macrophages prevented the cleavage of several previously described GP63 targets. These results point to the importance of inhibiting GP63 activity during the preparation of *Leishmania*-infected host cell lysates. In addition, the role(s) of GP63 in *Leishmania* pathogenesis must be reassessed in light of the findings reported in this study.

## RESULTS

### GP63-mediated proteolysis of several host cell proteins is a post-cell lysis event

Host cell PTPs, which play a central role in dampening the microbicidal functions in *Leishmania*-infected macrophages, are among reported GP63 substrates (15, 26-28). These PTPs include SHP-1, TC-PTP, PTP1B, and PTP-PEST and were shown to be activated through GP63-mediated proteolytic cleavage (21, 29). We thus examined and compared the fate of two representative PTPs, PTP-PEST and SHP-1, when cell lysates from infected BMM were prepared in the absence or presence of the zinc chelator 1,10-phenanthroline to efficiently inhibit the proteolytic activity of GP63 (23). To this end, BMM were infected with either WT, isogenic GP63-defective *Δgp63* mutants, or complemented *Δgp63*+*GP63 L. major* metacyclic promastigotes. At 2 h, 6 h, and 24 h post-infection, cell lysates were prepared in standard lysis buffer or in lysis buffer containing 10 mM 1,10-phenanthroline, and the levels of PTP-PEST and SHP-1 were assessed by Western blot analyses. As previously reported, both PTP-PEST and SHP-1 show important degradation/cleavage in lysates of BMM infected with either WT or *Δgp63*+*GP63* parasites prepared in standard lysis buffer (Figure 1A). In contrast, no cleavage of either PTPs was observed in lysates of BMM infected with the GP63-defective *Δgp63* mutant. Strikingly, cleavage of PTP-PEST was completely abrogated when infected cells were lysed in the presence of 10 mM 1,10-phenanthroline (Figure 1A). These results suggest that GP63-mediated cleavage of PTP-PEST observed in cell lysates prepared in standard lysis buffer is a post-cell lysis proteolytic event. To confirm this hypothesis, we mixed cell extracts from WT *L. major* promastigotes with lysates of BMM prepared in standard lysis buffer or in lysis buffer containing 2 or 10 mM 1,10-phenanthroline. As shown in Figure 1B, PTP-PEST was degraded when we mixed *L. major* promastigotes and BMM lysates prepared in standard lysis buffer. When cell extracts were prepared in lysis buffer containing either 2 or 10 mM 1,10-phenanthroline, we did not observed degradation of PTP-PEST (Figure 1B), consistent with the notion that GP63-mediated degradation of PTP-PEST occurred during cell processing. Of note, the previously reported GP63-mediated degradation of PTP-PEST was observed in *L. major*-infected fibroblasts (29). In the case of SHP-1, we observed a discrete cleavage product in macrophages infected with either WT or *Δgp63*+*GP63 L. major* metacyclic promastigotes (Figure 1A) when cell lysates were prepared in the presence of 10 mM 1,10-phenanthroline. This observation indicates that SHP-1 is a genuine GP63 substrate in infected BMM. Consistent with previous reports indicating that GP63-mediated cleavage/degradation of PTPs modulates their phosphatase activity (21, 29), we observed dephosphorylation of a subset of macrophages proteins following infection with either WT or *Δgp63*+*GP63 L. major* promastigotes (Figure 1C). However, dephosphorylation of these proteins was significantly attenuated when cell extracts were prepared in lysis buffer containing phosphatase inhibitors.

**Figure 1.**
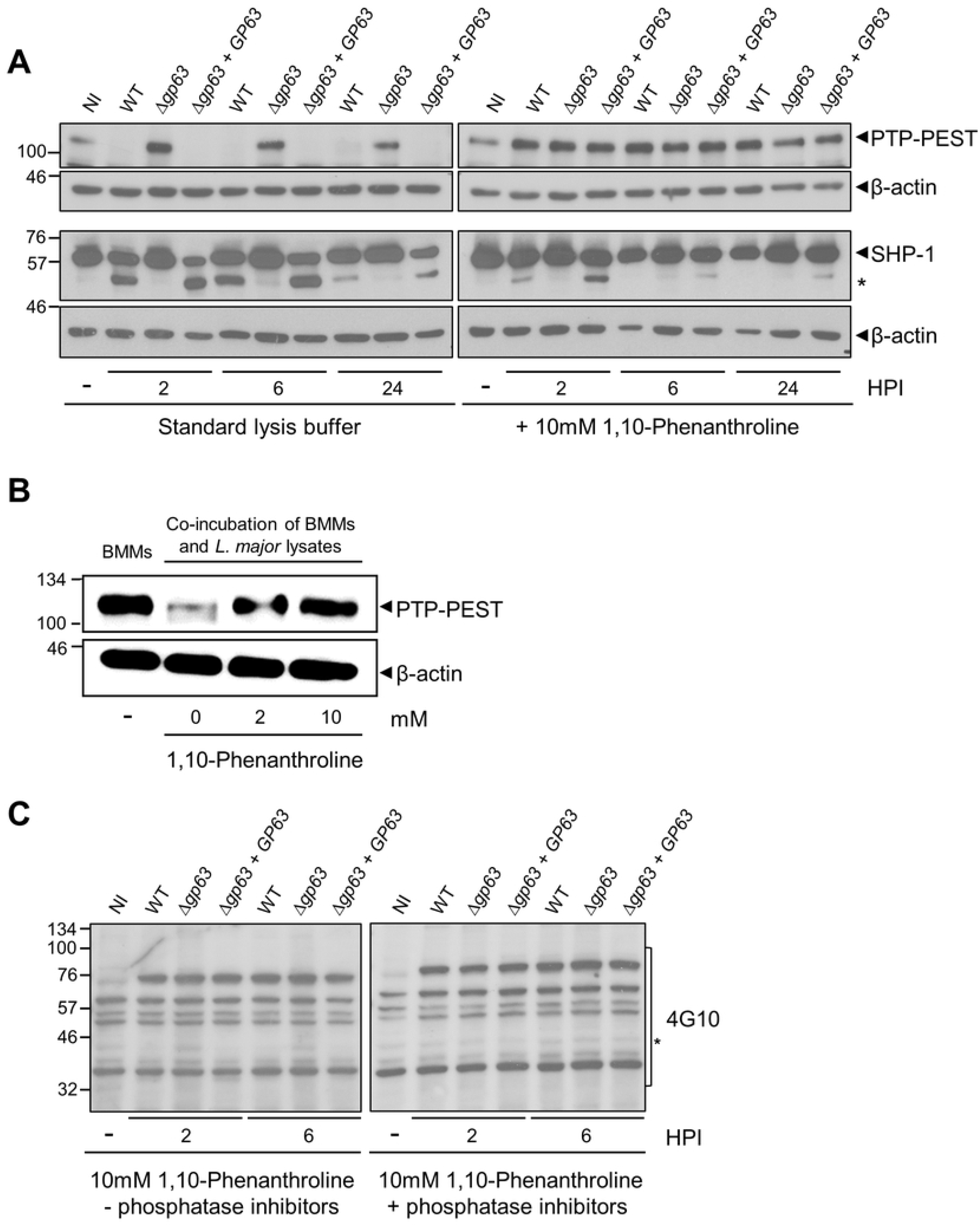
Re-assessment of host cell PTPs as GP63 substrates. A. Lysates of uninfected BMM (0 h post-infection) or BMM infected with either WT, *Δgp63*, or *Δgp63*+*GP63 L. major* metacyclic promastigotes were prepared at the indicated time points in standard lysis buffer or in lysis buffer containing 10 mM 1,10-phenanthroline. Levels and integrity of PTP-PEST and SHP-1 were assessed by Western blot analysis. β-actin was used a loading control. Asterisk denotes cleavage fragment. B. Lysates of uninfected BMM and of WT *L. major* promastigotes were prepared in in standard lysis buffer or in lysis buffer containing either 2 or 10 mM 1,10-phenanthroline. BMM and *L. major* promastigotes lysates were mixed for 15 min and the levels and integrity of PTP-PEST were assessed by Western blot analysis. β-actin was used a loading control. C. Lysates of uninfected BMM (0 h post-infection) or BMM infected WT, *Δgp63*, or *Δgp63*+*GP63 L. major* metacyclic promastigotes were prepared at the indicated time points in lysis buffer containing 10 mM 1,10-phenanthroline, in the absence or presence of phosphatase inhibitors (see *Materials and Methods*). Total protein tyrosine phosphorylation was assessed by Western blot analysis. Asterisk denotes dephosphorylated protein. Immunoblots shown are representative of at least four independent experiments. NI, non-infected BMM; HPI, hours post-infection.

These results prompted us to examine the fate of four additional randomly chosen GP63 published substrates: mTOR (30), the NLRP3 inflammasome (31), the transcription factor NF-κB subunit p65RelA (32), and the transcription factor c-Jun (33). As previously reported, at 2 h, 6 h, and 24 h post-infection, these proteins were degraded in a GP63-dependent manner when cell lysates were prepared in standard lysis buffer (Figure 2). In contrast, we did not detect degradation of these host proteins when lysates of infected BMM were prepared in the presence 10 mM 1,10-phenanthroline. These results suggest that similar to PTP-PEST, degradation of mTOR, NLRP3, NF-κB p65RelA, and c-Jun observed in infected BMM is a post-cell lysis proteolytic event. Given that both NF-κB, and c-Jun (AP-1) are transcription factors that mediate macrophage pro-inflammatory gene expression (34, 35), we measured the expression of *TNF, NOS2, IL6, IL1A*, and *IL1B* by RT-qPCR in BMM infected with either WT, *Δgp63*, or *Δgp63*+*GP63 L. major* metacyclic promastigotes. As shown in Figure 3, only *IL1A* expression was induced by *L. major* metacyclic promastigotes and its mRNA levels were not influenced by the presence of GP63. These results are consistent with the absence of detectable alterations of NF-κB and c-Jun in *L. major*-infected BMM and indicate that GP63 does not influence the expression of those pro-inflammatory genes during the infection of BMM with *L. major*.

**Figure 2.**
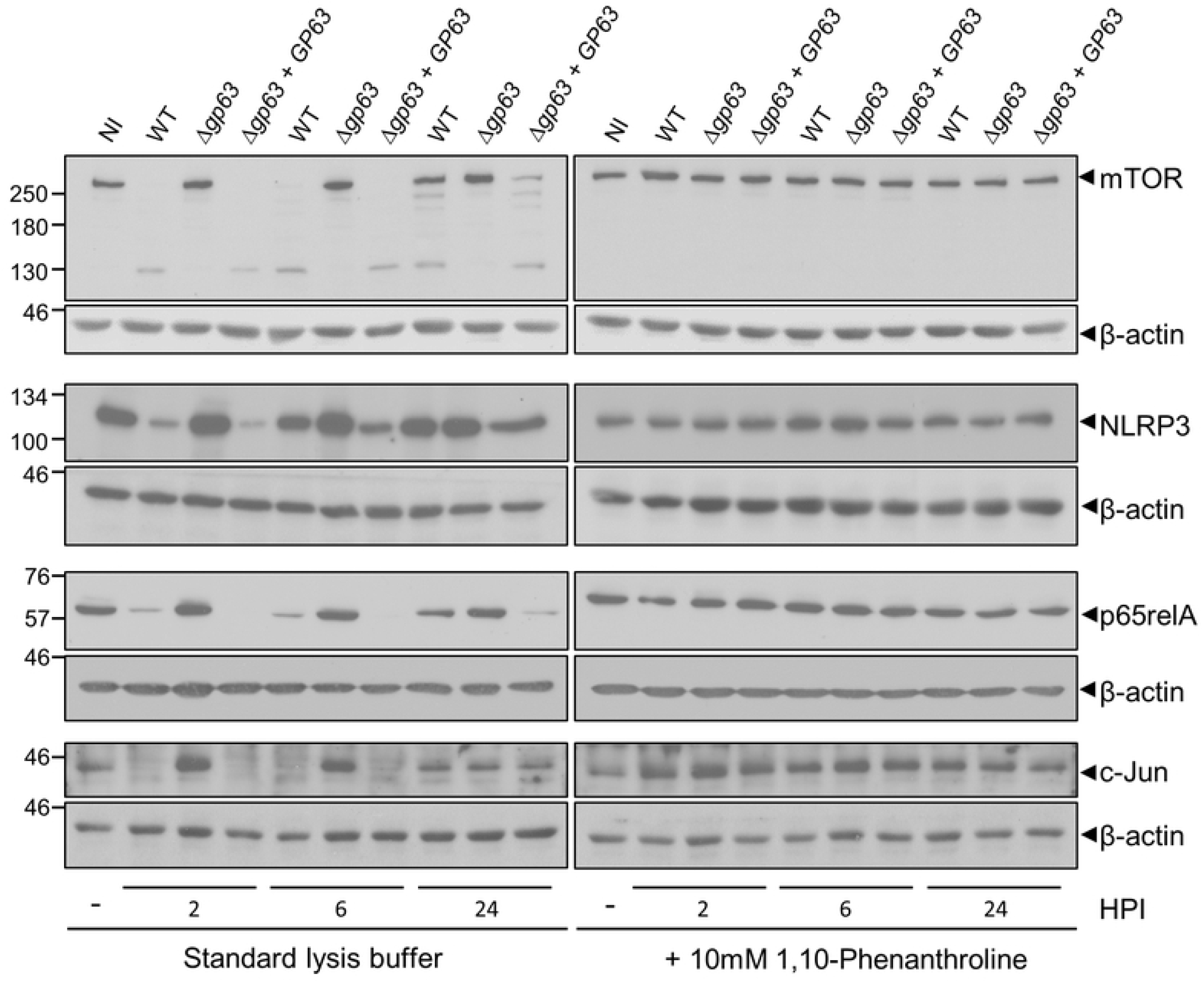
Degradation of mTOR, NLRP3, p65Rel, and c-Jun is prevented when GP63 activity is inhibited during cell lysate preparation. Lysates of uninfected BMM (0 h post-infection) or BMM infected with either WT, *Δgp63*, or *Δgp63*+*GP63 L. major* metacyclic promastigotes were prepared at the indicated time points in standard lysis buffer or in lysis buffer containing 10 mM 1,10-phenanthroline. Levels and integrity of mTOR, NLRP3, p65RelA, and c-Jun were assessed by Western blot analysis. β-actin was used a loading control. Immunoblots shown are representative of at least four independent experiments. NI, non-infected BMM; HPI, hours post-infection.

**Figure 3.**
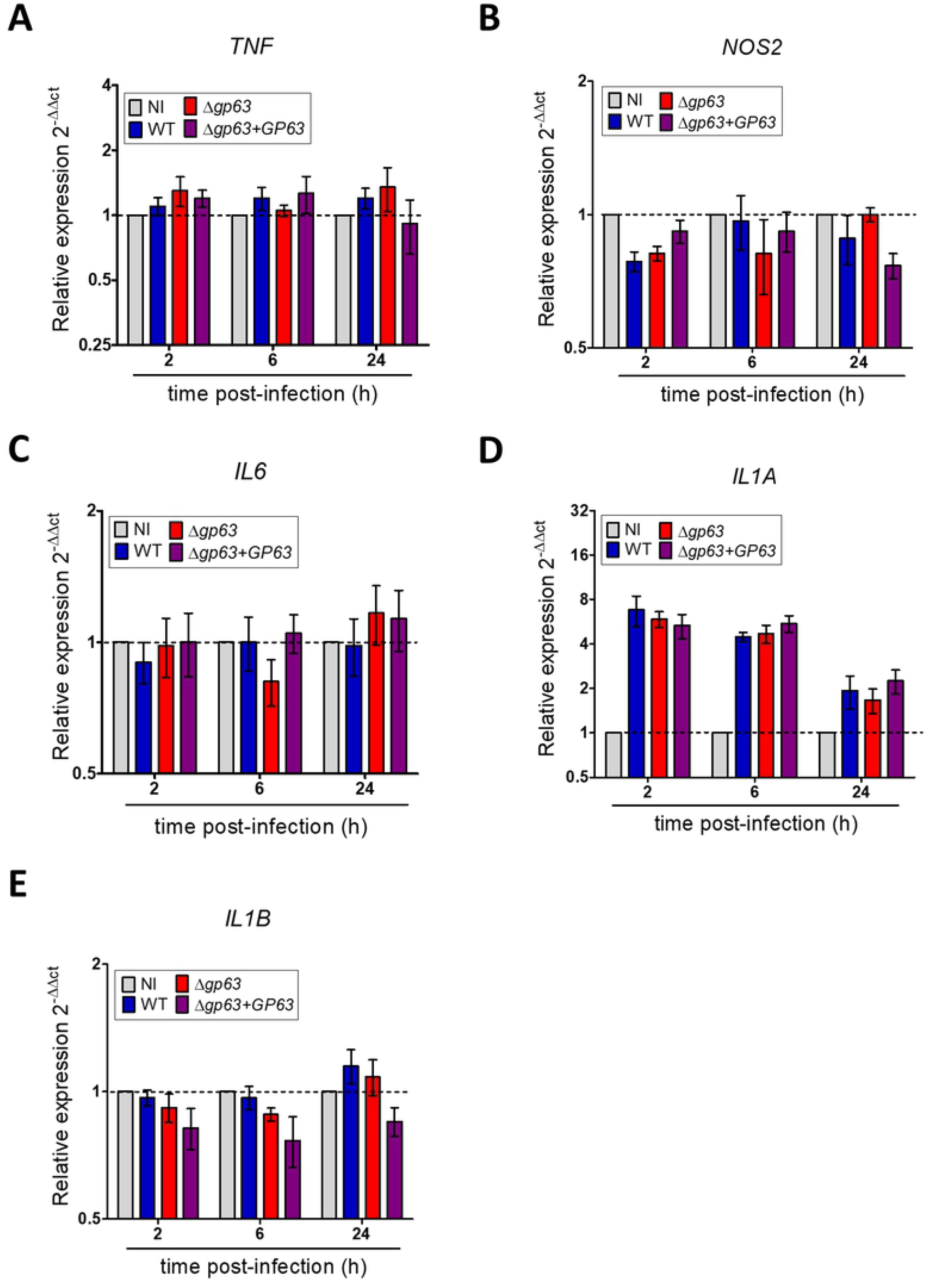
GP63 has no impact on the expression of macrophage pro-inflammatory genes. BMM were infected with either WT, *Δgp63*, or *Δgp63*+*GP63 L. major* metacyclic promastigotes and at 2 h, 6 h, and 24 h post infection the transcript levels for (A) *TNF*, (B) *NOS2*, (C) *IL6*, (D) *IL1A*, and (E) *IL1B* were analyzed by RT-qPCR. Data represent three independent experiments. The data were presented as means values ± SEM. NI, non-infected BMM.

### GP63 targets host cell regulators of membrane fusion

In a previous study, we identified the membrane fusion regulator Synaptotagmin XI (Syt XI) as a *bona fide* host cell GP63 substrate (36). In that study, we processed control and infected BMM in lysis buffer containing 2 mM 1,10-phenanthroline or alternatively, directly in 8 M urea prior to Western blot analyses (36). Urea was previously shown to effectively inhibit protease activity in lysates of *Chlamydia*-infected cells (37). To confirm these results, we infected BMM with either WT, *Δgp63*, or *Δgp63*+*GP63 L. major* metacyclic promastigotes for 1 h and 6 h prior to cell processing with either lysis buffer containing 2 mM 1,10-phenanthroline or by lysing cells in 8 M urea, as previously done. Figure 4A shows that Syt XI was degraded when cells were processed in the presence of 1,10-phenanthroline or in urea, indicating that proteolysis took place prior to cell processing. Together with the observation that Syt XI remained intact in extracts prepared from BMM infected with the *Δgp63* mutant (Figure 4A), our results confirmed that Syt XI is a genuine GP63 substrate. These results also indicate that addition of 2 mM 1,10-phenantroline in lysis buffer was sufficient to efficiently inhibit GP63 proteolytic activity. In addition to Syt XI, we previously reported that the SNAREs VAMP3 and VAMP8 were cleaved/degraded by GP63 in infected BMM (38). However, in that study, cell lysates were prepared in standard lysis buffer. We therefore compared lysates from infected BMM prepared in the absence or presence of 10 mM 1,10-phenanthroline at 2 h, 6 h, and 24 h post-infection. As shown in Figure 4B, no degradation of VAMP3 was detected when lysates of BMM infected with either WT or *Δgp63*+*GP63* were prepared in the presence of 1,10-phenanthroline. These results indicate that degradation of VAMP3 in *L. major*-infected BMM was a post-cell lysis event. For VAMP8, we detected a discrete cleavage in lysates of BMM infected with either WT or *Δgp63*+*GP63* prepared in the presence of 1,10-phenanthroline. In contrast, no cleavage of VAMP8 was observed in lysates of BMM infected with the GP63-defective *Δgp63* mutant (Figure 4B). To further illustrate the impact of GP63 on VAMP8, we analysed the presence of VAMP8 in infected BMM by confocal immunofluorescence microscopy at 6 h post-infection. As shown in Figure 4C, the intensity of labelling for VAMP8 was reduced in BMM infected either WT or *Δgp63*+*GP63 L. major* promastigotes compared to BMM that had internalized *Δgp63* parasites or to uninfected BMM. Together, these results are consistent with VAMP8 being a genuine GP63 host cell substrate.

**Figure 4.**
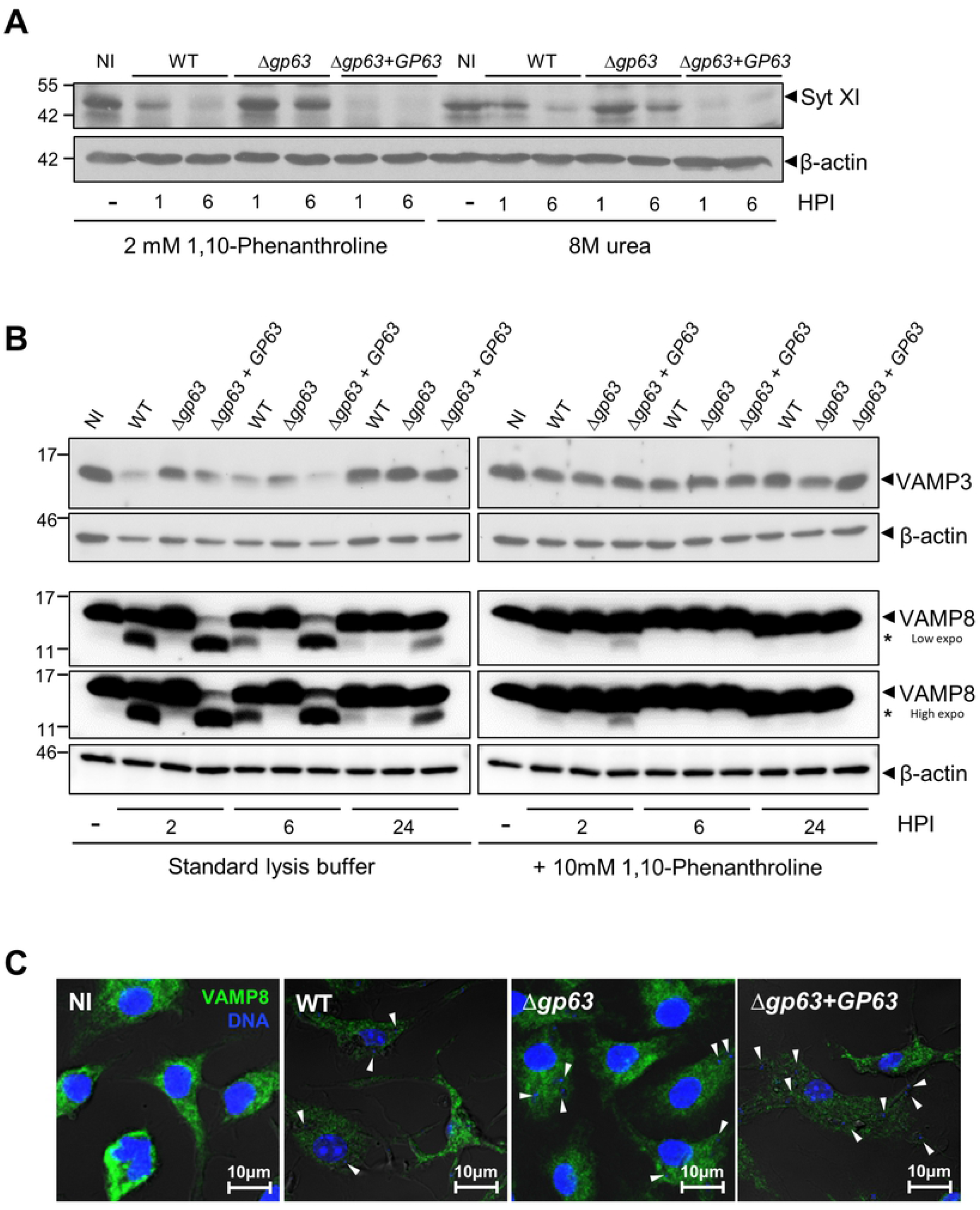
Syt XI and VAMP8 are *bona fide* GP63 substrates. A. Lysates of uninfected BMM (0 h post-infection) or BMM infected with either WT, *Δgp63*, or *Δgp63*+*GP63 L. major* metacyclic promastigotes were prepared at the indicated time points in lysis buffer containing 2 mM 1,10-phenanthroline or were lysed directly in 8 M urea. Levels and integrity of Syt XI were assessed by Western blot analysis. β-actin was used a loading control. B, C. Lysates of uninfected BMM (0 h post-infection) or BMM infected with either WT, *Δgp63*, or *Δgp63*+*GP63 L. major* metacyclic promastigotes were prepared at the indicated time points in standard lysis buffer or in lysis buffer containing 10 mM 1,10-phenanthroline. Levels and integrity of VAMP3 (B) and VAMP8 (C) were assessed by Western blot analysis. β-actin was used a loading control. Asterisk denotes cleavage fragments. Immunoblots shown are representative of at least four independent experiments. NI, non-infected BMM; HPI, hours post-infection. D. BMM were infected with either WT, *Δgp63*, or *Δgp63*+*GP63 L. major* metacyclic promastigotes for 6 h and the distribution of VAMP8 (green) was examined by immunofluorescence confocal microscopy. White arrowheads denote internalized parasites. These results are representative of three independent experiments. DNA is in blue; bar, 10 µM.

The SNARE Syntaxin-5 (Stx5) was previously shown to contribute to the expansion of communal parasitophorous vacuoles containing *L. amazonensis* (39) and to mediate the *Leishmania* effectors trafficking throughout infected cells (22). Given the role of this SNARE in the context of *Leishmania* infection, we sought to determine whether it is targeted by GP63 by comparing lysates from infected BMM prepared in the absence or presence of 10 mM 1,10-phenanthroline at 2 h, 6 h, and 24 h post-infection. Stx5 exists as a 35 kDa isoform (Stx5 short form or Stx5S) mainly present in the Golgi and a 42 kDa isoform (Stx5 long form or Stx5L) localized in the Golgi as well as in the ER (40). Similar to other host cell proteins assessed in the present study, both Stx5 isoforms were degraded in a GP63-dependent manner when cell lysates were prepared in standard lysis buffer (Figure 5A). In contrast, when cell lysates were prepared in the presence of 10 mM 1,10-phenanthroline, we observed a selective GP63-mediated cleavage of Stx5L whereas Stx5S remained intact. We further confirmed the selective degradation of Stx5L by confocal immunofluorescence microscopy. As shown in Figure 5B, the intensity of Stx5 staining was significantly reduced in the cytoplasm of BMM infected with either WT or *Δgp63*+*GP63 L. major* promastigotes, compared to the intensity of the Stx5 signal from the perinuclear structure (corresponding to the Golgi). These results are consistent with Stx5L being a *bona fide* GP63 host cell target. In addition, the selective cleavage of the ER-associated Stx5L and not the Golgi-associated Stx5S illustrates the notion that the action of GP63 within infected cells is compartmentalized.

**Figure 5.**
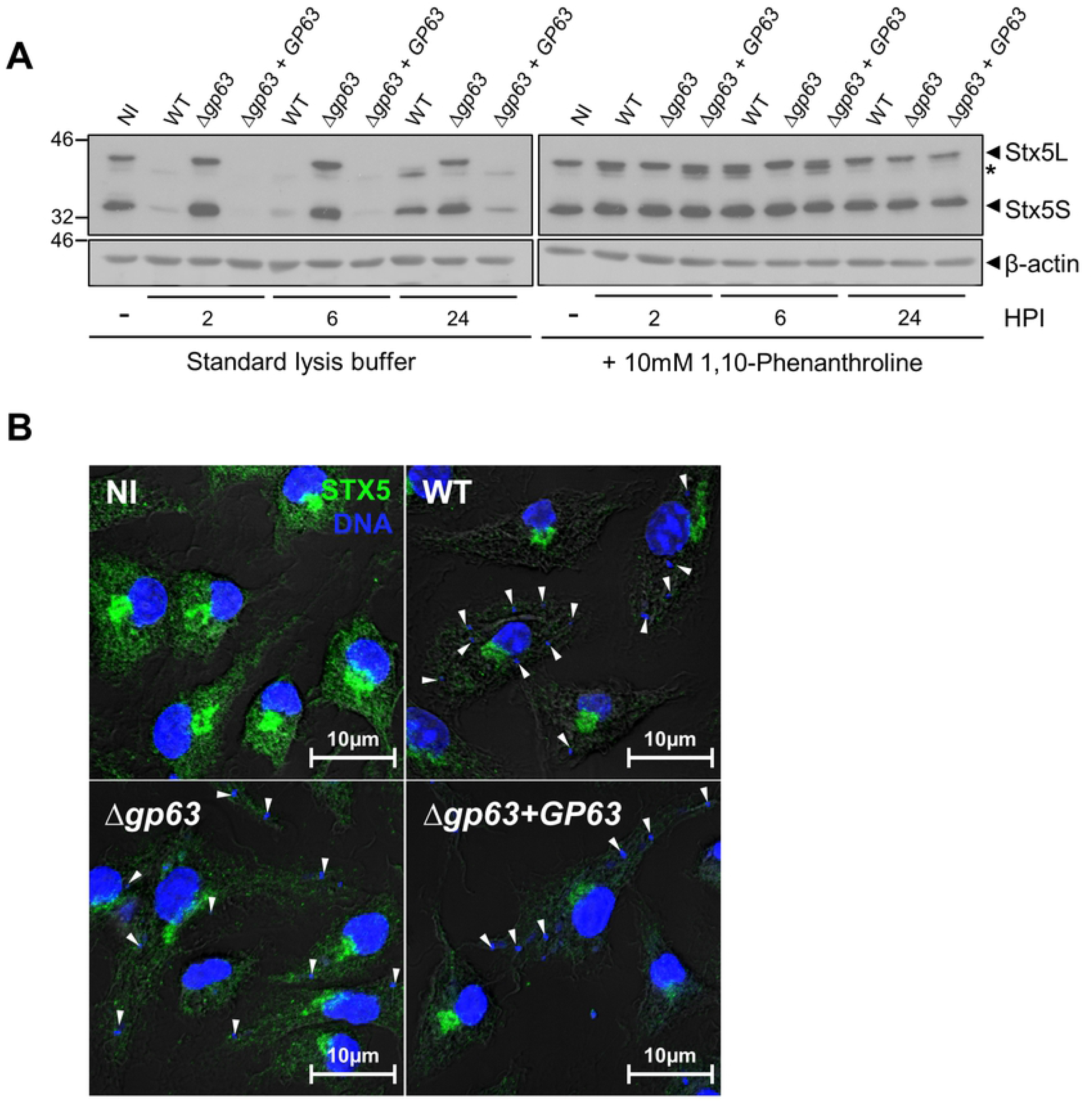
Stx5L is a genuine GP63 target. A. Lysates of uninfected BMM (0 h post-infection) or BMM infected with either WT, *Δgp63*, or *Δgp63*+*GP63 L. major* metacyclic promastigotes were prepared at the indicated time points in standard lysis buffer or in lysis buffer containing 10 mM 1,10-phenanthroline. Levels and integrity of Stx5L and Stx5S were assessed by Western blot analysis. β-actin was used a loading control. Asterisk denotes cleavage fragments. Immunoblots shown are representative of at least four independent experiments. NI, non-infected BMM; HPI, hours post-infection. B. BMM were infected with either WT, *Δgp63*, or *Δgp63*+*GP63 L. major* metacyclic promastigotes for 6 h and the distribution of VAMP8 (green) was examined by immunofluorescence confocal microscopy. White arrowheads denote internalized parasites. These results are representative of three independent experiments. DNA is in blue; bar, 10 µM.

## DISCUSSION

The abundant *Leishmania* zinc-metalloprotease GP63 is one of the pathogenicity factors that contribute to the colonization of mammalian hosts through the modulation of antimicrobial mechanisms and immune evasion (13, 15, 21, 30, 36, 38, 41). The main mode of action of GP63 resides in the cleavage/degradation of key host proteins that regulate processes involved in host defense against infection. Identification of GP63 substrates thus represents an essential step to understand how *Leishmania* establishes infection, replicates within mammalian hosts and evades immune processes. However, the powerful proteolytic activity of GP63 and its resistance to most protease inhibitors commonly used during the preparation of cell lysates (23) may have led to the identification of host proteins degraded by GP63 during cell processing. In this study, we have revisited the cleavage/degradation of ten host proteins previously described as GP63 substrates (21, 29-33, 36, 38). We found that presence of the metalloprotease inhibitor 1,10-phenanthroline during cell processing prevented the proteolysis of six published GP63 substrates evaluated in this study (PTP-PEST, mTOR, NLRP3, p65RelA, c-Jun, and VAMP3), indicating that these host cell proteins were degraded by GP63 during/after cell processing. We also provided evidence that SHP-1, Syt XI, VAMP8, and Stx5L are *bona fide* GP63 host cell substrates.

The availability of a genetically-defined *L. major* GP63 defective mutant (*Δgp63*) (13) and of transgenic GP63-altered *L. amazonensis* and *L. mexicana* (42, 43) was instrumental to unravel the role played by GP63 in the modulation of host phenotypes/responses associated to *Leishmania* infection (21, 30, 31, 33, 36, 38, 43-47). In addition, the availability of these tools allowed to further understand how *Leishmania* subverts and manipulates host cell processes to ensure its survival and replication, through the identification of several GP63 substrates (21, 30, 31, 33, 36, 38, 43-45). The finding that six of the previously identified GP63 substrates were not cleaved/degraded when cell lysates were prepared under appropriate conditions illustrates the importance of taking steps to effectively neutralize the activity of proteases present in cell lysates. Western blotting is widely used to assess protein levels, kinase and phosphatase activities, protein interaction, as well as post-translational modifications. However, there are limitations associated to this approach, since detection of discrete changes in a protein of interest may be hampered by several factors, including properties of the antibodies and sensitivity of the detection method (48). Thus, we cannot definitively rule out the possibility that discrete GP63-mediated cleavage events of host cell proteins investigated in the present study were below the detection limits of Western blot analysis. Alternative approaches such as immunofluorescence confocal microscopy could provide additional information on the potential impact of GP63 for those proteins for which we failed to detect cleavage/degradation by Western blot analyses. Hence, several other studies demonstrating the impact of GP63 on *bona fide* substrates related to signaling and nuclear structural proteins have been further supported by confocal microscopy observations. For instance, GP63-mediated cleavage/degradation related to PTP activation (e.g. SHP-1, PTP1B, nuclear PTP1C), nuclear isolated JUN/FOS family members in macrophage nuclei, and of several nucleoporins from nuclear membrane and nucleoplasm, also revealed the capacity of GP63 to rapidly reach the nuclear environment (21, 33, 45).

Out of the four *bona fide* GP63 substrates confirmed in this study, three are known regulators of phagolysosome biogenesis through the modulation of membrane fusion. Hence, SHP-1, VAMP8, and Syt XI control the phagosomal recruitment of the v-ATPase and of NOX2 (38, 49-52). In the case of Stx5, whereas little is known concerning its role in macrophages, previous work revealed that this SNARE contributes to the expansion of *L. amazonensis*-harboring communal parasitophorous vacuoles (39). Concerning Stx5S and Stx5L, current evidence indicates that each isoform interacts with distinct partners and plays specific functions (53-55). Of note, it was shown that Stx5L contributes to the organization of the ER through interactions with microtubules and CLIMP-63, an ER protein that regulates ER structure (54). Collectively, these results are consistent with the notion that interfering with the generation of a microbicidal phagolysosome through GP63-dependent proteolytic cleavage of regulators of membrane fusion may be part of the virulence strategy by *Leishmania* used to colonize host phagocytes (17).

In conclusion, our work illustrates the importance of preparing *Leishmania*-infected cell lysates under conditions that efficiently prevent proteolytic degradation during cell processing. In this regard, our results along with our recent report on *L. donovani*-inducible mTOR activity (56) indicate that the zinc-metalloprotease inhibitor 1,10-phenanthroline should be systematically included in lysis buffers used to prepare cell lysates. Alternatively, direct cell lysis in 8 M urea (37) or in sample loading buffer (for end-point application such as Western blot analyses) are an effective way of preventing post-cell lysis proteolytic events. This is particularly true in the context of studies involving protein analyses, including immunoblotting, identification of protein binding partners, and large-scale proteomics. Our work also raises the need to re-assess the role of GP63 in *Leishmania* pathogenesis.

## MATERIALS AND METHODS

### Ethics statement

Animal work was performed as stipulated by protocols 2112-01 and 2110-04, which were approved by the *Comité Institutionnel de Protection des Animaux* of the INRS-Centre Armand-Frappier Santé Biotechnologie. These protocols respect procedures on animal practice stipulated by the Canadian Council on Animal Care (CCAC), described in the Guide to the Care and Use of Experimental Animals.

### Bone marrow derived macrophages

Bone marrow-derived macrophages (BMM) were prepared as previously described (57). Briefly, marrow was extracted from femurs and tibias of 8 to 12 week-old male and female mice and differentiated for 7 days into BMM in Dulbecco’s Modified Eagle’s Medium with glutamine (DMEM, Thermo Fisher Scientific), containing 10% heat-inactivated, fetal bovine serum (Wisent), 10 mM HEPES pH 7.4, penicillin (100 I.U./mL), streptomycin (100 μg/mL) and supplemented with 15% (v/v) L929 cell-conditioned medium as a source of colony-stimulating factor-1 (CSF-1), in a 37°C incubator with 5% CO_2_. BMM were made quiescent by culturing them in DMEM without CSF-1 for 16 h prior to infection.

### Parasite cultures

Promastigotes of *L. major* Seidman (MHOM/SN/74/Seidman) NIH clone A2 and the isogenic Δ*gp63* mutants and their complemented counterparts Δ*gp63*+*GP63* (13) were cultured at 26° C in *Leishmania* medium (M199 [Sigma] supplemented with 10% v/v heat-inactivated fetal bovine serum (FBS), 40 mM HEPES pH 7.4, 100 µM hypoxanthine, 5 µM hemin, 3 µM biopterin, 1 µM biotin, and antibiotics). *L. major* Δ*gp63*+*GP63* promastigotes were cultured in medium supplemented with 100 μg/ml G418 (Life Technologies) (36). The *Leishmania* strains used in this study were passaged in mice to maintain their virulence and were kept in culture for a maximum of six passages. For infections, metacyclic promastigotes were enriched from late stationary-phase cultures using Ficoll gradients, as described (58).

### Infections

Metacyclic promastigotes were opsonized with C5-deficient serum from DBA/2 mice, resuspended in complete DMEM and fed to macrophages (MOI of 10:1) as previously described (36). To optimize contact, the multiwell plates were spun 2 min at 300 *g* in a Sorvall RT7 centrifuge. BMM were then incubated at 37°C and after 3 hours of incubation, non-internalized parasites were washed 3X with warm Hank’s Balanced Salt Solution.

### Cell lysis and Western blotting

Cells were washed 3X with cold PBS containing 1 mM of Na_3_VO_4_ (Sigma), with or without 5 mM of 1,10-phenanthroline (Sigma). The cells were lysed on ice in 100 µL of cold 1% NP-40, 50 mM Tris-HCl (pH 7.5), 150 mM NaCl, 1 mM EDTA (pH 8), 1.5 mM EGTA, 1 mM Na_3_VO_4_, 50 mM NaF, 10 mM Na_4_P_2_O_7_, and a mixture of protease and phosphatase inhibitors (Roche). When specified, 1,10-phenanthroline (Sigma) was added to the lysis buffer at the indicated concentration. Cell lysates were collected on ice with a cell scraper, transferred to a microtube, and stored at -70°C. Prior to use, cell lysates were centrifuged for 20 min to remove insoluble matter. Protein quantification was done using the Pierce® BCA Protein Assay Kit (Thermo Scientific) according to the protocol provided by the manufacturer. After protein quantification, the solution containing 20 µg of protein was boiled (100°C) for 5 min in Laemmli buffer and migrated in SDS-PAGE gels. Proteins were transferred onto Hybond-ECL membranes (Amersham Biosciences), blocked for 1 h in TBS 1X-0.1% Tween 20 containing 5% BSA (bovine serum albumin from BioShop), incubated with primary antibodies (diluted in TBS 1X-0.1% Tween 20 containing 5% BSA) overnight at 4°C, and then with the according HRP-conjugated secondary antibodies for 1 h at room temperature. Membranes were then incubated in ECL (GE Healthcare) and immunodetection was achieved via chemiluminescence.

### Co-incubation of cell lysates

BMM and *L. major* promastigotes were collected and lysed independently on ice with standard lysis buffer or with lysis buffer supplemented with either 2 or 10 mM of 1,10-phenanthroline. Lysates from BMM and *L. major* were then co-incubated on ice for 15 min and stored at - 70°C prior to Western blot analyses.

### Antibodies

Rabbit polyclonal antibodies against Syntaxin-5, VAMP3, and VAMP8 were obtained from Synaptic Systems; rabbit polyclonal against β-actin, p65RelA, and mTOR from Cell Signaling; rabbit polyclonal anti-c-Jun from Santa Cruz Biotechnology; mouse monoclonal anti-SHP-1 from Abcam; mouse monoclonal antibody anti-NLRP3 from AdipoGen Life Sciences. The rabbit anti-Syt XI polyclonal antibody (59) was kindly provided by Dr Mitsunori Fukuda (Tohoku University). The rabbit polyclonal antibodies anti-PTP-PEST 2530 (60) were kindly provided by Dr Michel L. Tremblay (McGill University). The secondary HRP-conjugated antibodies anti-mouse and anti-rabbit were obtained from Sigma Aldrich. The secondary antibody anti-rabbit linked with AlexaFluor 488 used for immunofluorescence was from Invitrogen-Molecular probes.

### Immunofluorescence

Cells on coverslips were fixed with 3.8% paraformaldehyde (Thermo Scientifc) for 30 min at 37 ° C. NH_4_Cl were used to eliminate free radicals in the cells for 10 min. The cells were permeabilized for 5 min with a 0.5% Triton X-100. The samples were subsequently blocked for 1 h at 37°C with PBS containing 5% FBS. Primary antibodies were diluted 1/200 in PBS containing 5% FBS and incubated for 2 h at 37°C. Then, the coverslips were incubated for 1 h at 37°C in a solution containing the secondary antibodies bound to AlexaFluor 488 (diluted 1: 500) and DAPI (1: 20000, Molecular Probes, D3571) in PBS + 5% FBS. The coverslips were washed three times with 1x PBS between each step. The coverslips were mounted on glass slides (Fisher) with Fluoromount-G (Southern Biotechnology Associates, 010001). Images of the cells were captured using the 63X objective of an LSM780 confocal microscope (Carl Zeiss Microimaging). Control stainings confirmed no cross-reactivity or background. Images were taken in sequential scanning mode via ZEN software. Images were analyzed with Icy Software from the Institut Pasteur (61). The statistical analysis of the data was done with the GraphPad Prism 5 software. The mean of the data ±SEM of three independent experiment was analyzed. Data were considered statistically significant when *** P<0,001, ** P<0.01, according to a unidirectional ANOVA with a Tuckey multiple comparison test.

### Quantitative PCR analysis

Total BMM RNA was isolated using RNeasy mini kit (QIAGEN), according to the manufacturer’s protocol. Five hundred nanograms of RNA were reverse transcribed using the iScript cDNA Synthesis kit (Bio-Rad). Real-time quantitative PCR (RT-qPCR) reactions, performed on at least three independent biological replicates, were run in duplicate by using iTaq Universal SYBR® Green Supermix (Bio-Rad) on a Stratagene mx3005p real-time PCR system using 10ng cDNA. Gene expression changes were analyzed using the comparative CT method (ΔΔCT) (62). Relative mRNA amounts were normalized to the Rps29 gene and expressed as fold increase to non-infected controls. The following primers were used. *Rps29*: 5’-CACCCAGCAGACAGACAAACTG-3’ (F) and 5’-GCACTCATCGTAGCGTTCCA-3’ (R); *IL1A*: 5’-CAA ACT GAT GAA GCT CGT CA-3’ (F) and 5’-TCT CCT TGA GCG CTC ACG AA-3’ (R); *IL1B*: 5’-AAC CTG CTG GTG TGT GAC GTT C-3’ (F) and 5’-CAG CAC GAG GCT TTT TTG TTG T-3’ (R); *IL6*: 5’ACAACCACGGCCTTCCCTACTT-3’ (F) and 5’-CACGATTTCCCAGAGAACATGTG-3’ (R); *NOS2*: 5’-CAAGATGCGTGGAAACTACC-3’ (F) and 5’-TTGAGAATGGATGCGAAGG-3’ (R); and *TNF*: 5’-GAC GTG GAA GTG GCA GAA GAG-3’ and 5’-TGC CAC AAG CAG GAA TGA GA-3’ (R).

## ACKNOWLEDGMENTS

We thank Dr. Mitsunori Fukuda (Tohoku University) for the kind gift of the rabbit anti-Syt XI, Dr. Michel L. Trembay (McGill University) for the kind gift and the rabbit anti-PTP-PEST, Dr. W. Robert McMaster (University of British Columbia) for kindly providing the *L. major* lines, and Jessy Tremblay for assistance in immunfluorescence experiments. This work was supported by Canadian Institutes of Health Research grant PJT-156416 to AD. AD holds the Canada Research Chair on the Biology of intracellular parasitism. MJ holds a salary award Junior 2 from the Fonds de recherche du Québec - Santé. MMGV was the recipient of a Master Student Scholarship from the Fondation Armand-Frappier. AMB was the recipient of a Natural Science and Engineering Research Council of Canada Undergraduate Student Research Award. The funders had no role in study design, data collection and analysis, decision to publish, or preparation of the manuscript.

## AUTHOR CONTRIBUTIONS

Conceptualization, M.-M.G.-V., C.M., and A.D.; Methodology, M.-M.G.-V, C.M., A.-M.B., and A.D.; Formal Analysis, M.-M.G.-V, C.M., A.-M.B., and A.D.; Investigation, M.-M.G.-V., C.M., A.-M.B.; Resources, A.D.; Writing - Original Draft, M.-M.G.-V. and A.D.; Writing – Review and Editing, A.D., M.J., C.M., MO; Visualization, M.-M.G.-V., C.M., A.-M.B., A.D.; Supervision, A.D.; Funding Acquisition, A.D.

